# A behavioral comparison of hybrid virtual reality and real-world object lifting

**DOI:** 10.64898/2026.04.13.718283

**Authors:** Catherine Anne Sager, Jackson Zenti, Michelle Marneweck

**Author notes:** Correspondence: Michelle Marneweck (, 1240 University of Oregon Eugene OR 97403).

## Abstract

Object manipulation requires anticipatory force control and online force adjustments to accommodate variability in digit position. Appropriate force scaling is learned rapidly, but repeated manipulation can interfere with updating control when object dynamics change. Sensory feedback and prior experience support object manipulation, yet how proprioception contributes remains poorly understood. Hybrid virtual reality (VR), in which participants manipulate a physical object while viewing tracked virtual representations of the object and their digits, allows controlled visual-proprioceptive discrepancies to be introduced to examine visual and proprioceptive contributions to control. However, replacing direct viewing with virtual feedback could itself affect these control processes. We therefore compared manipulation of the same physical object under direct vision and hybrid-VR without experimentally imposed visual-proprioceptive offsets. Fifteen healthy young adults lifted a visually symmetric inverted T-shaped object with a concealed asymmetric mass distribution while preventing tilt toward the weighted side. After either one or five repeated lifts, the mass distribution switched to the opposite side. In both environments, anticipatory torque increased with repeated lifts, and five preceding lifts produced greater post-switch error than one, consistent with repetition-induced anterograde interference. Digit position and lift-force differences were negatively correlated during later pre-switch trials in both environments. We detected no significant differences between environments in the learning rate of anticipatory force control, position-force coordination, or switch-related interference. These findings characterize object manipulation under no-offset hybrid-VR and provide a baseline for future studies using controlled visual-proprioceptive discrepancies to examine how these signals support anticipatory control, learning, and online adjustment.

**NEW AND NOTEWORTHY:** Hybrid virtual reality (VR), combining physical object interaction with virtual visual feedback, showed rapid anticipatory force learning, position-force coordination during later pre-switch trials, and repetition-induced anterograde interference. No significant differences from direct vision were observed on these measures, providing a baseline for future studies introducing visuo-proprioceptive discrepancies to examine how these sensory signals contribute to object control.

## 1. INTRODUCTION

Object manipulation requires precise coordination of digit position and force (1–5). Before lift onset, digit forces are scaled according to expected object properties, such as mass and mass distribution (1,6). When these properties are uncertain, anticipatory force scaling can be learned rapidly, often within only a few trials (6,7). When object dynamics change, anticipatory force control must be updated, for example, to generate an appropriate compensatory torque following a switch in mass distribution. This update is not immediate. Instead, early post-switch behavior often reflects anterograde interference (8,9) in which force and torque outputs remain biased toward the pre-switch condition. This carryover is magnified when the pre-switch condition is extensively repeated (*i.e*., repetition-induced anterograde interference) (7,10). In addition to anticipatory control and learning, skilled object manipulation requires online force adjustments to accommodate trial-to-trial variability in digit position (4,5,11).

Anticipatory control, learning, and online adjustments during object manipulation are supported by sensory feedback and sensorimotor memories acquired through prior object interactions (1,11–14). Visual information provides cues about object properties such as size (15) and shape (16) and can cue the associated sensorimotor memory (15,17). Even when properties such as mass distribution are not visually salient, vision remains important. For example, when an object’s mass distribution is visually concealed, vision at and after grasp contact facilitates learning following changes in object dynamics, consistent with a role in monitoring performance and updating sensorimotor memories (18). It also supports efficient coordination of grip and lift forces, with faster force generation despite greater trial-to-trial variability in digit placement (18). Although sensory feedback (18–24) and prior experience (4,6–8,10,12,13) are well-established supports of anticipatory and online force control, the contribution of proprioception remains poorly understood.

Studies of clinical deafferentation have informed our understanding of proprioceptive contributions to motor control (25–30), including object manipulation (19,20). However, behavior following profound, chronic sensory loss also reflects long-term compensatory strategies, making the effects of sensory loss difficult to distinguish from adaptation to it (21,30). Hybrid virtual reality (VR), in which participants physically interact with a real object while viewing a tracked virtual representation, provides an experimental approach to examining the relative contributions of vision and proprioception to object manipulation. It allows controlled discrepancies between these signals to be introduced by manipulating visual information while retaining proprioceptive input from physical interaction. Controlled visuo-proprioceptive discrepancies have been used in reaching to examine the relative contributions of these sensory signals (31,32). However, given the contribution of vision to object manipulation, replacing direct viewing with a tracked virtual representation could itself affect anticipatory control, learning, and online adjustments. Thus, before using hybrid-VR to dissociate visual and proprioceptive influences, it is necessary to establish whether the mode of visual presentation itself alters object manipulation.

Accordingly, we compared manipulation of the same physical object under direct vision and hybrid-VR, in which participants viewed a tracked virtual representation of the object and their digits without experimentally imposed visual-proprioceptive offsets. We examined early learning of anticipatory force control, trial-to-trial coordination of digit positions and forces, and susceptibility to repetition-induced anterograde interference. Participants lifted a visually symmetric inverted T-shaped object with a concealed asymmetric mass distribution while preventing tilt toward the weighted side. The present study does not dissociate the contributions of visual and proprioceptive information to object manipulation. Instead, characterizing performance under no-offset hybrid-VR provides a baseline for future studies using controlled visual-proprioceptive discrepancies to examine their relative contributions to anticipatory control, learning, and online adjustments.

## 2. MATERIALS AND METHODS

Participants completed a well-established asymmetric-load object lifting task (4–7,9,10,12,33–36) in real-world and hybrid-VR environments. In this task, participants reached, grasped, and lifted an inverted T-shaped object with a concealed off-centered mass while preventing it from tilting toward the weighted side. Successful performance required anticipatory force control, achieved by generating a compensatory torque at lift onset (Figure 1A).

**Figure 1:**
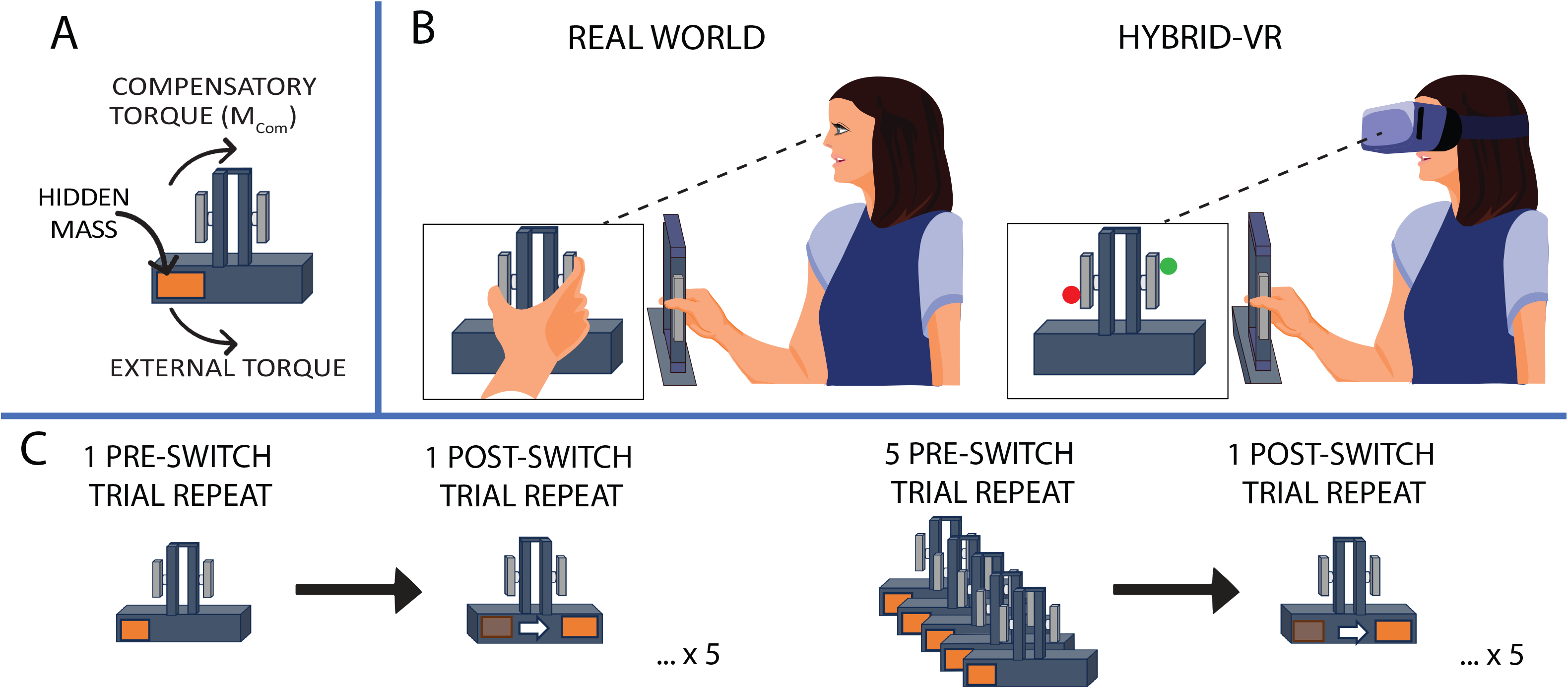
Object and Task Design. A) Inverted T-shaped object with a concealed off-center mass. Participants (N = 15) had the task goal of minimizing tilt by generating a compensatory torque (M_Com_) that opposes the object’s external torque. B) Lifting the inverted T-shaped object with the right index finger and thumb with their fingertips in the real world (left) and hybrid virtual reality (VR) (right). In the real world, participants saw the object and their grasping hand/digits throughout the reach, grasp, and object lift (left box); in hybrid-VR, participants saw a visual representation of the object in VR and their grasping digits (thumb: red marble; index finger: green marble) throughout the reach, grasp, and lift (right box). C) We experimentally manipulated the number of pre-switch trial repeats (1 or 5) with the CoM on a given side (*left*, in the example) before switching the CoM to the other side (*right*, in the example).

### 2.1 Participants

Fifteen healthy right-handed young adults (median age: 23, range: 18-35, standard deviation = 2.8; 10 females) participated in this study. The study and all its procedures were approved by the University of Oregon Institutional Review Board, and all participants gave written informed consent.

### 2.2 Apparatus

The T-object was fabricated with aluminum rods and plates. The object’s vertical column (height: 10.2 cm; width: 1.0 cm; depth: 3.4 cm) had smooth elongated grasp surfaces (height: 7.0 cm; width: 0.3 cm; depth: 4.1 cm; between grasp distance: 4.6 cm) with 400 grain sandpaper attached on either side of the grasp surfaces. A lead cylinder (height: 4.5 cm; diameter: 2.4 cm; mass: 215 g) was concealed in the horizontal base (height: 3.8 cm; width: 32.3 cm; depth: 3.8 cm). The total mass of the object was 650 g with an external torque of 250 Nmm. A three-camera motion tracking system (Precision Point Tracking System; WorldViz) with a frame rate of 150 Hz (camera resolution: 1,280 × 1,024 VGA) was used to track the vertical height of the object. The system’s spatial accuracy within a 3 × 3 × 3 m volume was ≤ 1 mm. Two near-infrared LED markers were affixed securely to the covers on the horizontal base of the object with the midpoint between the two markers (vertical column of the object) used as a reference point to monitor the object’s vertical position relative to the table. Grip forces and torque that were applied to the grip surfaces of the object were recorded at a frequency of 500 Hz through force/torque transducers (Mini27 Titanium, ATI Industrial Automation, NC). These force transducers were attached between each grip surface and the vertical column of the T-shaped object. Grasp surfaces were coupled to the transducers such that forces and torques measured at each sensor reflected the forces applied by the thumb and index finger at the grasp surfaces. Grip force, load forces, and torque, with resolutions of 0.03 N, 0.015 N, and 0.375 Nmm were measured by the transducers.

For the VR task, participants wore an HTC VIVE headset (2160 × 1200 resolution, PenTile OLED display; HTC VIVE, Xindian District, New Taipei City, Taiwan) equipped with integrated headphones (HTC VIVE Tracker (3.0) Developer Guideline ver. 1.1, 2021) and sampled at a rate of 120 Hz. A 3D virtual environment was rendered using the Unity3D game engine (v2018.4.10f1; Unity Technologies, San Francisco, USA) with the SteamVR plugin. The HTC VIVE tracking system used SteamVR infrared tracking to record the real-time position and orientation of two near-infrared LED markers affixed to the inverted T-shaped object, as well as two additional markers placed on the participant’s right index finger and thumb. In the VR display, the finger markers were represented as marble-sized spheres (green for the index finger and red for the thumb; Figure 1B). The custom VR environment included a white floor, a virtual desk with a tape box indicating where to replace the object, a keyboard button, and the inverted T-shaped object used for the lifting task. Auditory cues were presented through the headset’s headphones.

### 2.3 Experimental Procedures

In both real-world and hybrid-VR environments, participants were seated in a chair at a comfortable height such that their right forearm and upper arm made a right angle while resting on the table. They were instructed to keep their left arm resting in their lap. At the start of each trial, participants pressed a keyboard button (home button) with their right index finger, 20 cm from the T-shaped object that they would interact with throughout the task. To test for anticipatory force control, a “left” or “right” audio cue informed them which side of the inverted T-shaped object would be heavier. A go beep occurred 1 s after the left/right side audio cue that instructed participants to reach to lift the object without tilting it. Participants were informed that they could grasp the object anywhere along the elongated grasp surfaces with their fingertips. The object was lifted to a height of 11 cm. A second beep (2.50 s after the go cue) (5,18,37,38) instructed participants to set the object back down in a taped square on the table and return their right index finger to its start position on the keyboard button. If the object rolled more than 5 degrees to either side, an error tone would sound. Participants were told to complete both tasks at a natural pace and to attempt to lift the object without tilting it to the weighted side at all times. Mean reach duration, calculated from button-up time to when participants applied more than 0.2 N of horizontal (grip) force on the T-shaped object, was 1.03 s (SD = 0.26) in hybrid-VR and 0.95 s (SD = 0.27) in the real-world environment, leaving approximately 1.5 s in both environments to lift the object before the set-down cue; reach duration did not differ between environments (*p* > .05).

To examine whether real-world and hybrid-VR object lifting show similar effects of anticipatory force control learning and repetition-induced anterograde interference, we manipulated the number of trial repeats in the object-use task before implementing a switch in the object’s center of mass (CoM). Participants completed 10 independent blocks in total: five blocks comprising 1 pre-switch trial followed by 1 post-switch trial and five blocks comprising 5 pre-switch trials followed by 1 post-switch trial (Figure 1C), yielding 40 trials in total (30 pre-switch and 10 post-switch lifts). Within each block, the five pre-switch trials involved the same CoM location, followed by one post-switch trial with the opposite CoM. After the post-switch trial, the asymmetric mass was returned to the participant’s assigned pre-switch CoM configuration before the start of the next block. Thus, the first pre-switch trial of Blocks 2–5 followed a between-block change back to the assigned pre-switch CoM, whereas subsequent pre-switch trials within each block followed a trial with the same CoM location. Participants were assigned either 1) right pre-switch and left post-switch trials or 2) left pre-switch and right post-switch trials. The pre-switch CoM location (left or right) was counterbalanced across participants. Both real-world and hybrid-VR environments were completed by the same participants (within-subjects design) in two separate sessions (48 hours apart). The order of environments (real-world *vs*. hybrid-VR) and pre-switch trial condition (1 vs. 5) were counterbalanced between subjects.

### 2.4 Data Processing

Kinetic and kinematic preprocessing were conducted in Python (version 3.12) using custom scripts with NumPy and signal-processing routines, including peak detection and low-pass filtering. Data were filtered using a fourth-order low-pass Butterworth filter with a cutoff frequency of 5 Hz.

The following primary outcome measures of anticipatory control were quantified at lift onset, defined as the point at which the object was displaced upward by at least 5 mm and had a continuously increasing vertical position over 20 consecutive samples.

1. Grip force mean (GF_mean_) at lift onset is the instantaneous average in grip force of each digit in Newtons (N). This was calculated using a numerical averaging method:

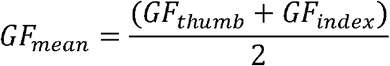
2. Lift force difference (LF_diff_) at lift onset is the difference in the tangential component of the force produced by each digit (N).

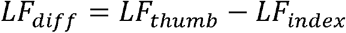 Positive values indicate greater thumb than index-finger lift force, whereas negative values indicate greater index-finger than thumb lift force. Larger differences indicate a more asymmetric lift force sharing pattern, whereas a zero value indicates a symmetric lift force sharing pattern.
3. Center of pressure (COP) at lift onset is the measure of digit position defined as the point of contact of each digit on the grip surface relative to the center of the transducer (in mm). This was computed using the formula:

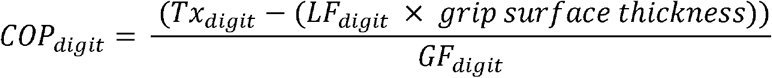 where T*x*_digit_ is the digit torque in the frontal plane (Nmm). The center of pressure difference (COP_diff_) between thumb and index finger placement was used to identify grip configuration:

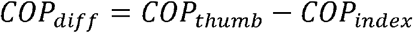

where positive values indicate higher thumb than index finger COP and negative values indicate higher index finger than thumb COP. Larger differences indicate a more asymmetric, non-collinear grip configuration, whereas a zero value indicates a symmetric, collinear grip configuration.
4. Compensatory moment or torque (M_Com_) at lift onset is the anticipatory torque generated by the digits measured in Newton millimeters (Nmm) to counter the external torque of the object. This was computed using the formula:

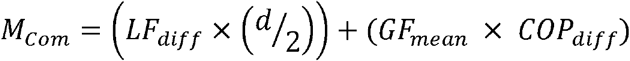 where *d* is the width between the two grip surfaces (4.6 cm). A positive M_Com_ indicated a clockwise moment, and a negative M_Com_ indicated a counterclockwise moment.

For participants who began the task with the CoM on the right, M_Com,_ LF_diff_, and COP_diff_ values were multiplied by -1 to avoid the statistical complication caused by different signs of M_Com_ when manipulating an object with a left *vs*. a right CoM (7,10,18). Prior research (10) confirmed that a starting CoM position (left vs. right) did not significantly influence pre- and post-switch lift performance, justifying pooling of trials across CoM positions.

### 2.5 Data Analyses

To assess within-CoM learning of anticipatory force control, we focused on the 5 pre-switch lifts in each environment (real world, hybrid-VR). To characterize the nonlinear time course of early adaptation, we used a nonlinear mixed-effects exponential learning model:

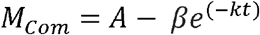

where denotes trial within each pre-switch sequence (*t* = 0, . . ., 4), *A* represents the asymptotic (plateau) M_Com_, *β* represents the difference between the asymptote and predicted M_Com_ on the first trial (*t* = 0), and *k* represents the learning-rate constant. The model included participant-specific random effects for *A* (*A*~|*Subject*). The fixed-effects interaction structure for *k* is as follows:

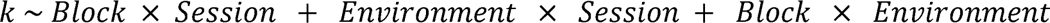

This specification includes the main effects of Environment, Block, and Session, together with Block × Session, Environment × Session, and Block × Environment interactions; a model that also included the three-way interaction did not converge. Block was treated as a categorical factor with five levels. Fixed effects for *A* and *β* were intercept-only. Models were fit by maximum likelihood. Factors were effect-coded (sum-to-zero contrasts). Fixed effects were evaluated with marginal *F*-tests. Parameter estimates and contrasts are reported with 95% confidence intervals (CIs). With Environment and Session linked within participants in this counterbalanced crossover, the Environment × Session interaction corresponds to the order effect. To test whether Environment or Session affected the level rather than the rate of learning, these terms were added to *A* and evaluated with a likelihood-ratio test; the resulting Environment × Session contrast on *A*, which compares the two order groups, was evaluated with 13 degrees of freedom.

To characterize early learning in individual M_Com_ components (COP_diff_, LF_diff_, and GF_mean_), we conducted separate two-way repeated-measures ANOVAs with Trial (1, 2, 3, 4, and 5) and Environment (real-world, hybrid-VR) as within-subject factors. Block and Session were not included in these component-level ANOVAs because the nonlinear mixed-effects analysis of M_Com_ learning did not identify significant effects of Block or Session on learning rate or asymptote. Post-hoc Tukey comparisons were performed to examine differences between individual trials. M_Com_ comprises torque contributions from lift force (*LF_diff_* × *d2* and digit position (*GFmean* × *COPdiff*). To further characterize the strategy used to generate M_Com_, we calculated the relative contribution of the COP-dependent torque component. The COP-dependent contribution (R_COP_) was calculated as:

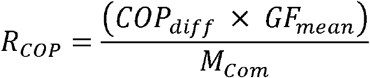

Values between 0 and 1 indicate that the components contributed in the same direction as the net M_Com_, whereas values greater than 1 or less than 0 indicate that the torque components acted in opposing directions. A three-way repeated-measures ANOVA was performed to determine whether R_COP_ differed across Trials (1, 2, 3, 4, 5), Blocks (1, 2, 3, 4, 5), and Environment (real-world, hybrid-VR). Session was not included because the nonlinear mixed-effects analysis of M_Com_ learning did not identify effects of Session on learning rate or asymptote.

To assess trial-by-trial coordination between digit positioning and digit lift force, we examined the relationship between COP_diff_ and LF_diff_ across the final three pre-switch lifts of each block (15 trials total) for each participant in each Environment (real-world, hybrid-VR). Post-switch trials were not included. We chose to restrict our principal position-force analysis to the later, more stable trials, allowing time for participants to adapt to each CoM reset and establish a position-force coordination policy; during the early stages of this adaptation, participants are likely less able to adjust force appropriately to digit position (confirmed by secondary analyses below). For each participant, Pearson’s correlation coefficients (*r*) were calculated between COP_diff_ and LF_diff_, and then Fisher *r*-to-*z* transformed for statistical testing. Participants’ individual Fisher *z* values were compared between real-world and hybrid-VR environments using a paired-samples *t*-test to understand whether the strength of the relationship between digit positioning and digit lift force differed across environments. Group-level mean *r* values were reported to describe the overall magnitude of these correlations. We performed secondary sensitivity analyses using 1) pre-switch lifts from blocks 3–5 (15 trials total) and 2) five pre-switch lifts from the five-trial pre-switch condition (for a total of 25 trials) for each participant in each Environment (real-world, hybrid-VR). Correlation coefficients and confidence intervals were calculated using the same approach described above.

To examine learning transfer between CoM conditions, we performed a mixed-design repeated-measures ANOVA on post-switch M_Com_ with pre-switch Repetition (1, 5), Environment (real-world, hybrid-VR), and Block (1, 2, 3, 4, and 5), as within-subjects factors. We also included Environment Order (real-world first, hybrid-VR first) as a between-subject factor to account for potential order effects. Because Environment and Session are linked within participants in this counterbalanced crossover, the Environment × Environment Order interaction in this analysis corresponds to the Session effect. Post-hoc Tukey comparisons were performed to examine differences between individual blocks.

Statistical analyses were performed in R (version 4.6.0; R Foundation for Statistical Computing, Vienna, Austria) using RStudio (version 2026.04.0+526; Posit, Boston, MA, USA). Nonlinear mixed-effects models were fit, and 95% CIs for their parameters obtained, with the nlme package (version 3.1-169), repeated-measures ANOVAs were conducted with the afex package (version 1.5-1), and estimated marginal means, contrasts, and post-hoc comparisons, with their 95% CIs, were obtained with the emmeans package (version 2.0.3). Greenhouse–Geisser corrections were applied where estimable. One trial from a single participant with missing force data was excluded from all relevant analyses; this participant was excluded from the R_COP_ analysis, where each design cell contains a single trial.

## 3. RESULTS

### 3.1 No Significant Effect of Environment on Anticipatory Torque Learning

Figure 2 shows early learning-related changes in anticipatory torque control (M_Com_) and its contributors (COP_diff_, LF_diff_, GF_mean_) across pre-switch trials in both real-world and hybrid-VR environments. M_Com_ improved across the first five pre-switch trials (β = 111 Nmm, 95% CI [95, 128]), approaching an asymptote of 176 Nmm (95% CI [153, 199]; full model results in Supplemental Table S1). The learning rate did not differ significantly between environments or sessions, there were no significant effects involving Block, and no interactions (Supplemental Tables S1–S2). Allowing the asymptote to vary by Environment and Session did not improve model fit, χ^2^(3) = 3.85, *p* = .278, with the between-participant SD being 39 Nmm for the asymptote.

**Figure 2:**
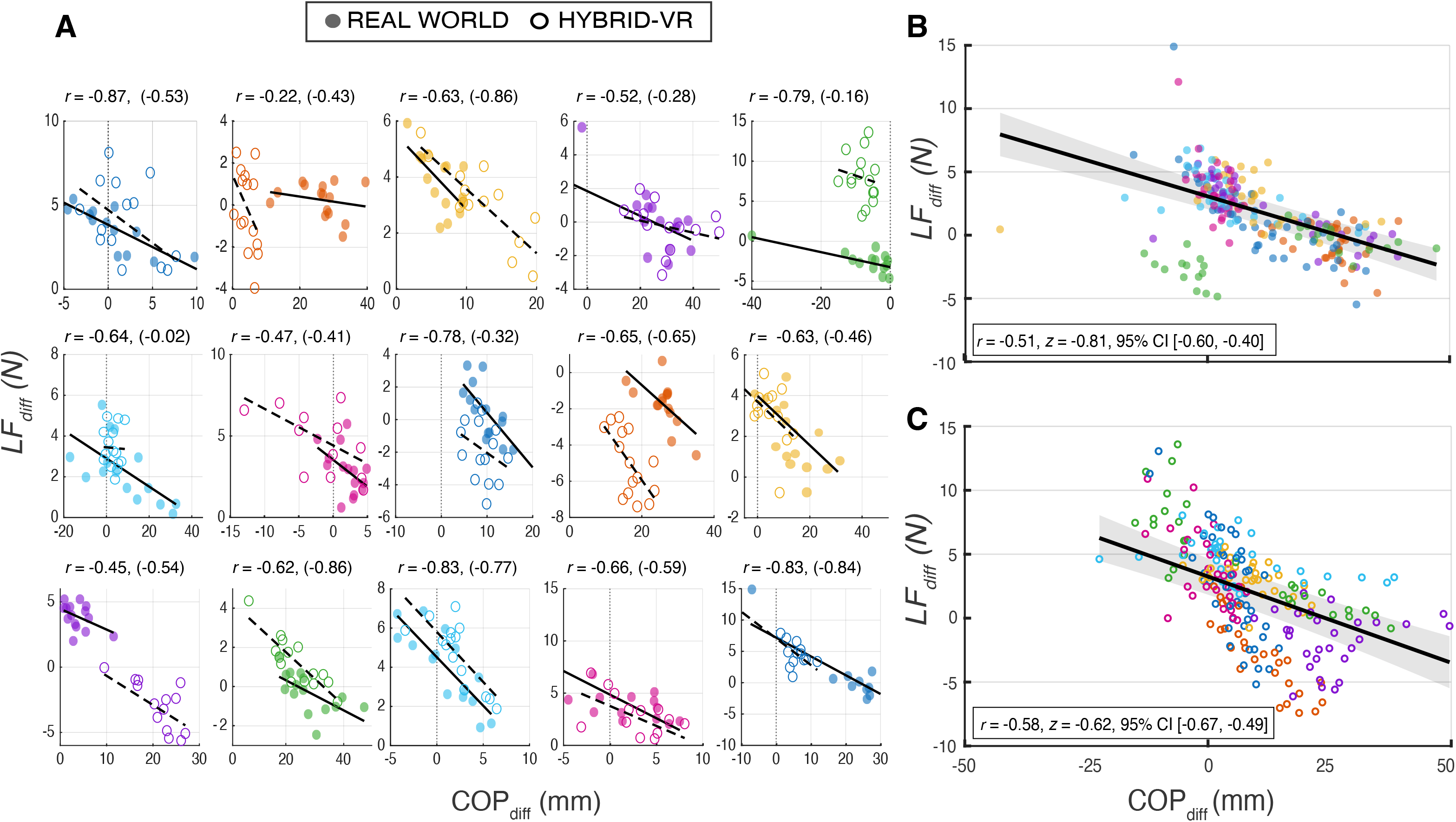
Learning Rate of Anticipatory Torque Control in Real-World and Hybrid Virtual Reality (VR) Object Manipulation. Left) Group (N = 15) mean compensatory moment (M_Com_) aggregated across five blocks of pre-switch trials in both real-world (purple) and hybrid-VR (yellow) environments. Increasing experience with a given center of mass led to improved anticipatory torque control in both environments. Right) Components of M_Com_ [center of pressure difference (COP_diff_), lift force difference (LF_diff_), and mean grip force (GF_mean_)] on five pre-switch trials collapsed across blocks showed no significant Environment effects or interactions. COP_diff_ and LF_diff_ showed significant trial-dependent changes, whereas GF_mean_ remained stable across early learning. (* *p* < .05, ** *p* < .01, *** *p* < .001). Error bars represent ± 1 SD.

COP_diff_ showed a significant main effect of Trial (*F*(3.14, 43.97) = 10.68, *p* < .001, η_p_² = .43), no significant main effect of Environment (*F*(1, 14) = 0.91, *p* = .356, η_p_² = .06), and no significant Trial × Environment interaction (*F*(2.58, 36.19) = 1.01, *p* = .389, η_p_² = .07). Tukey-adjusted pairwise comparisons indicated that COP was significantly greater on Trial 4 (*p* = .002) and Trial 5 (*p* = .005) than Trial 1 and Trial 4 (*p* = .007) and Trial 5 (*p* = .018) than Trial 2. No other pairwise comparisons were significant. Similarly, LF_diff_ showed a significant main effect of Trial (*F*(2.76, 38.62) = 17.23, *p* < .001, η_p_² = .55), no significant main effect of Environment (*F*(1, 14) = 0.01, *p* = .914, η_p_² < .01), and no significant Trial × Environment interaction (*F*(2.92, 40.83) = 1.31, *p* = .302, η_p_² = .09). Tukey-adjusted pairwise comparisons indicated that LF was significantly greater on Trials 2 (*p* < .001), 3 (*p* < .001), 4 (*p* = .004), and 5 (*p* < .001) than on Trial 1. In contrast, GF_mean_ showed no significant main effect of Trial (*F*(1.99, 27.91) = 1.04, *p* = .368, η_p_² = .07) or Environment (*F*(1, 14) = 3.97, *p* = .066, η_p_² = .22). The Trial × Environment interaction was also not significant (*F*(2.13, 29.87) = 0.28, *p* = .775, η_p_² = .02). For R_COP_, there were no significant main effects of Trial, *F*(1.38, 17.93) = 0.14, *p* = .792, η_p_² = .01; Environment, *F*(1, 13) = 0.06, *p* = .818, η_p_² < .01; or Block, *F*(2.59, 33.63) = 1.29, *p* = .292, η_p_² = .09. The Trial × Environment interaction, *F*(1.77, 23.03) = 0.73, *p* = .478, η_p_² = .05; Trial × Block interaction, *F*(16, 208) = 1.67, *p* = .054, η_p_² = .11; Environment × Block interaction, *F*(2.35, 30.58) = 1.56, *p* = .224, η_p_² = .11; and Trial × Environment × Block interaction, *F*(16, 208) = 1.29, *p* = .207, η_p_² = .09, were also nonsignificant (Supplemental Figure S1).

### 3.2 Trial-By-Trial Variation in Digit Positions and Lift Force Partitioning in Real World and Hybrid-VR Manipulation

To assess whether the hybrid-VR object lifting task exhibits trial-to-trial control processes similar to those observed during real-world manipulation, we quantified and compared the relationship between LF_diff_ and COP_diff_ across pre-switch lifts for each of these environments. Figure 3A shows individual participants’ trial-by-trial correlations between LF_diff_ and COP_diff_ for the last three trials of each block for both real-world and hybrid-VR environments. Group mean correlations when the final three trials of each block were included (15 total trials) showed significant negative LF_diff_–COP_diff_ correlations in both real-world (mean *r* = −0.51, 95% CI [−0.61, −0.40]; Figure 3B) and hybrid-VR (mean *r* = −0.58, 95% CI [−0.67, −0.49]; Figure 3C). Importantly, there was no significant statistical difference of the LF_diff_–COP_diff_ relationship between these environments (*p* > .05).

**Figure 3:**
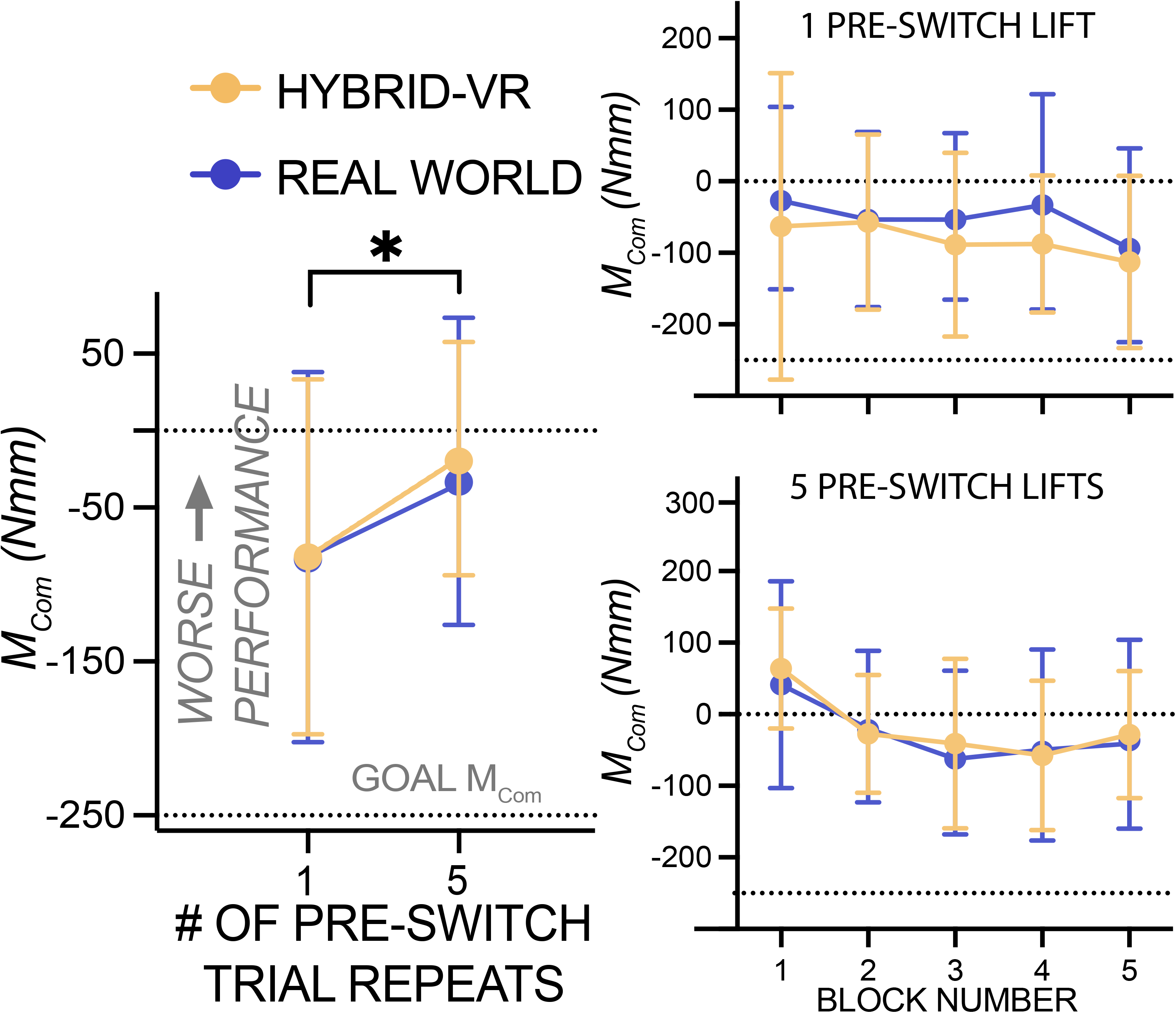
Trial-by-Trial Digit Position and Lift Force Partitioning Scatterplots for Real World and Hybrid Virtual Reality (VR) Manipulation. A) Individual participants’ (N = 15) trial-by-trial scatterplots and Pearson correlation coefficients (*r*) (hybrid-VR in parentheses) showing the relationship between lift force difference (LF_diff_) and center-of-pressure difference (COP_diff_) during object lifting in both real-world (closed circles) and hybrid-VR (open circles) environments using the last three trials of each block (Trials 3–5; 15 trials per participant). Different axis scales were used across participants for visualization purposes. Each subplot depicts data from a single participant, with colors identifying individuals. B) Group-level real-world scatterplot showing the relationship between LF_diff_ and COP_diff_ using the last three trials of each block (Trials 3–5; 15 trials per participant). C) Group-level hybrid-VR scatterplot showing the relationship between LF_diff_ and COP_diff_ using the last three trials of each block (Trials 3–5; 15 trials per participant). In both B and C, points are colored by participant, and the fitted regression line with its 95% confidence interval (CI) is shown.

In secondary analyses, we examined the relationship between COP_diff_ and LF_diff_ using the final three blocks and using all 25 pre-switch trials. Similar to when the last three trials of each block were used, when only trials from the final three blocks were included (15 total trials), significant negative correlations were observed in both environments (real-world: *r* = −0.42, 95% CI [-0.53, -0.31]; hybrid-VR: *r* = −0.51, 95% CI [−0.60, −0.41]). When all 25 pre-switch trials were used, group mean correlations were variable across participants in both real-world (mean *r* = −0.38, 95% CI [−0.70, 0.11]) and hybrid-VR (mean *r* = −0.44, 95% CI [−0.78, 0.13]), with confidence intervals spanning zero in both cases. Importantly, there was no significant statistical difference of the LF_diff_–COP_diff_ relationship between these environments (*p* > .05).

### 3.3 Presence of Repetition-Induced Anterograde Interference in the Lift Component of the Real World and Hybrid-VR Manipulation Tasks

Figure 4 shows that repetition-induced anterograde interference was magnified in post-switch trials as the number of pre-switch repetitions increased in both real-world and hybrid-VR environments. Results showed significant main effects of Repetition (*F*(1,13) = 7.60, *p* = .016, η_p_² = .36) and Block (*F*(2.45, 31.87) = 7.00, *p* = .002, η_p_² = .35). Post-hoc comparisons showed that Block 1 differed from 3 (*p* = .030), 4 (*p* = .001), and 5 (*p* = .044). Blocks 1-2 and 2-5 did not differ from each other (all *p*’s > .05). No significant main effects of Environment, Environment Order, or interactions involving Repetition, Environment, Environment Order, and Block were detected (all *p*’s > .05).

**Figure 4:**
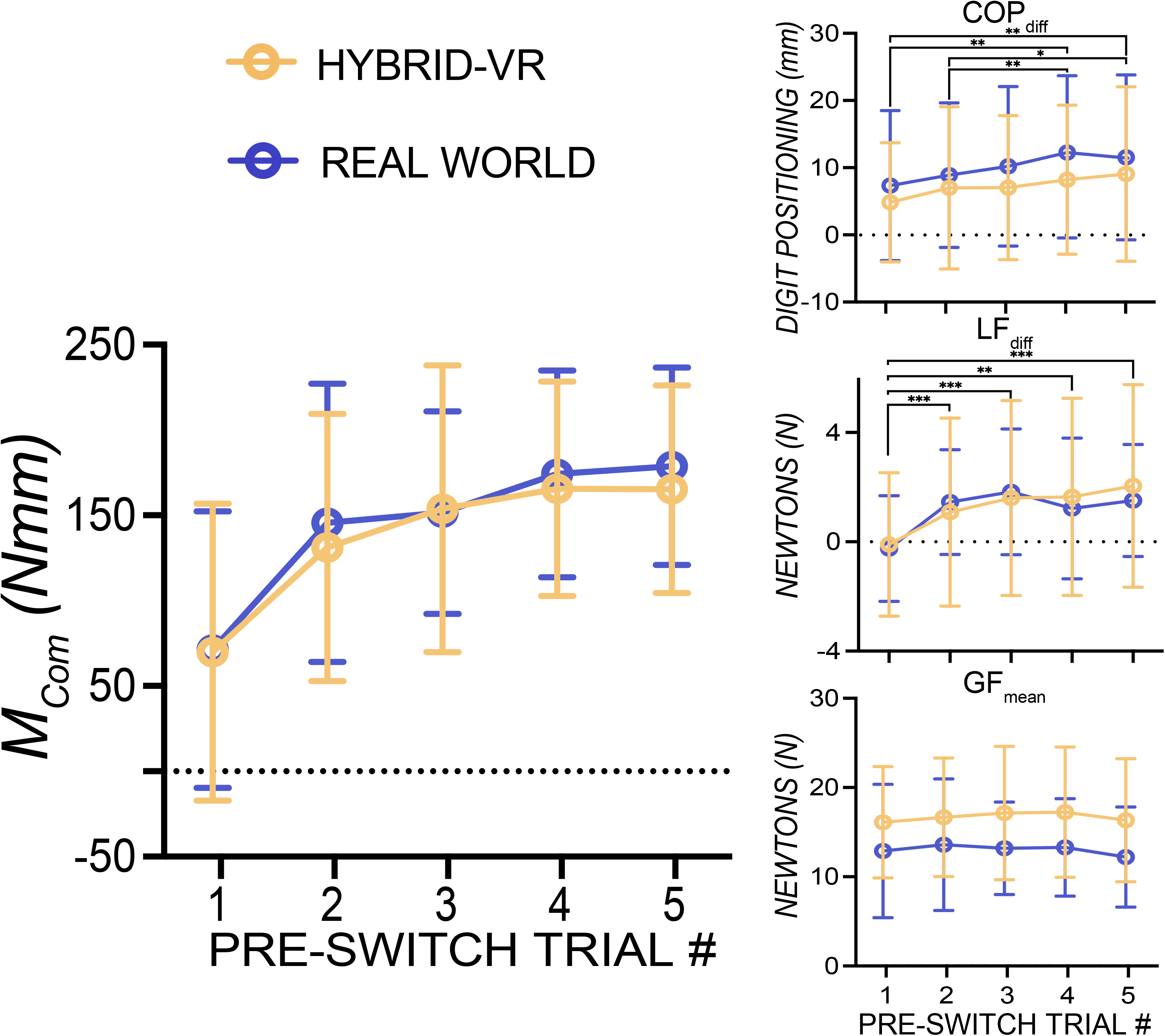
Repetition-Induced Anterograde Interference Present in Both Real World and Hybrid Virtual Reality (VR) Manipulation. Left) Group (N = 15) mean post-switch compensatory moment (M_Com_) following one and five pre-switch trials in the real-world (purple) and hybrid-VR (yellow) environments (collapsed across blocks). More positive values indicate poorer performance relative to the goal M_Com_ of −250 Nmm (marked with a horizontal dotted line). Five pre-switch lifts produced significantly greater post-switch errors than one pre-switch lift, demonstrating repetition-induced anterograde interference. Right) Group mean post-switch M_Com_ across Blocks 1-5 following one pre-switch lift (top) and five pre-switch lifts (bottom), shown separately for the real-world and the hybrid-VR environments. Anterograde interference decreased with blocks, with a significant difference between Block 1 and Blocks 3-5, whereas Blocks 1-2 and 2-5 did not differ from each other. No significant main effects of Environment or Environment Order and no interactions involving Environment, Repetition, Block, or Environment Order were detected. Asterisks indicate statistical significance (* *p* < .05). Error bars represent ± 1 SD.

## 4 DISCUSSION

In this study, we tested whether a hybrid-VR environment elicits core features of dexterous object manipulation observed in the real world. Specifically, we compared hybrid-VR with real-world physical object manipulation in learning of anticipatory force control, coordinated trial-to-trial variability in digit position and force, and susceptibility to repetition-induced anterograde interference. Participants performed an asymmetric object lifting task both in a real-world setting and in a hybrid-VR environment in which they manipulated a physical object while viewing it virtually. Consistent with our hypotheses, we detected no significant differences between environments in the learning rate of anticipatory control, trial-to-trial coupling between digit position and force, or the magnitude and persistence of repetition-induced anterograde interference, but these nonsignificant effects do not establish equivalence between environments.

Participants rapidly learned to generate an anticipatory torque to minimize object roll in both environments, with no significant differences in learning rate. This pattern is consistent with prior real-world studies showing that anticipatory torque is adjusted within the first few lifts of an object with an asymmetric mass distribution (1,6,17,39). Torque components also showed distinct trial-dependent modulation, with changes in lift-force partitioning evident earlier in the sequence than changes in digit position, while mean grip force and relative COP-dependent torque contribution remained stable. Critically, none of these patterns differed between environments, indicating that virtual visual feedback did not detectably alter either the overall learning of anticipatory torque or the component adjustments supporting it.

The trial-to-trial relationship between digit position and lift-force partitioning also did not differ significantly between environments. Prior real-world studies have reported moderate-to-strong negative relationships between digit position and lift force, indicating that participants adjust force to compensate for trial-to-trial variability in digit placement (4,5,18,40). In our primary analysis, which included the final three pre-switch trials of each block, position-force correlations were moderate and negative in both real-world and hybrid-VR environments and were similar in magnitude to those reported previously (4,5,18). We also conducted two secondary analyses to examine how the trial structure influenced this relationship. When analyses were restricted to the final three blocks, correlations remained similar in magnitude to those reported previously. In contrast, when all 25 pre-switch trials were included, correlations were smaller, and confidence intervals included zero. This reduction may reflect the inclusion of early trials following each CoM reset. Previous studies generally examined longer sequences of trials with the same CoM (e.g., 8+ trials) following practice or initial adaptation, whereas our trials were embedded within shorter five-trial blocks with repeated CoM switches. Because each block ended with the opposite CoM, the first pre-switch trial of Blocks 2–5 followed a lift performed with the opposite CoM location. Anticipatory control on these trials may therefore still have been influenced by the immediately preceding object dynamics, weakening the trial-to-trial relationship between digit position and lift-force partitioning. Thus, the position-force relationship was most evident when analyses emphasized later experience within or across blocks, with correlation magnitudes comparable to those reported previously. Across these analyses, there was no statistically significant difference between real-world and hybrid-VR environments.

Repetition-induced anterograde interference was evident in both real-world and hybrid-VR environments: post-switch error was greater after five than after one pre-switch lift and was larger in Block 1 than in blocks 3-5. Although block-collapsed torque remained in the direction required by the new mass distribution, after five pre-switch lifts, mean torque in Block 1 remained in the previously required direction, indicating negative transfer. The stronger interference with greater pre-switch exposure (7,8,10) and its attenuation with repeated switching (8,36) are consistent with prior work. Critically, neither the repetition effect nor the block-related change differed significantly between environments.

The present study introduced no visuo-proprioceptive discrepancies and therefore did not manipulate proprioception. Instead, it tested whether replacing direct vision with tracked virtual visual feedback altered established features of anticipatory and online object control. Because the CoM was visually concealed, participants could not determine the direction of the required compensatory torque from object geometry alone and instead had access to the auditory cue and sensorimotor memories acquired from previous lifts. Even when properties such as mass distribution are not visually salient, vision remains important, facilitating learning following changes in object dynamics, consistent with a role in monitoring performance and updating sensorimotor memories (18). It also supports efficient coordination of grip and lift forces, with faster force generation despite greater trial-to-trial variability in digit placement (18). Nevertheless, we detected no significant effect of replacing direct vision with virtual visual feedback on any of the anticipatory and online control measures examined here. Prior work suggests that congruent visual cues from shape and mass distribution may be more useful for anticipatory torque planning (41,42) than incongruent cues, in which object geometry does not provide a valid cue to an asymmetric mass distribution despite explicit knowledge of the CoM (43). A future comparison between real-world and hybrid-VR manipulation using an object with visually informative and congruent properties would provide an additional informative stress-test of whether replacing direct vision with virtual visual feedback affects control.

These findings provide a baseline for future studies introducing controlled visual-proprioceptive discrepancies to examine the relative contributions of visual and proprioceptive information during object manipulation. Proprioceptive information contributes to sensing the position and movement of body parts relative to one another and to the sense of muscle force (44). It may therefore contribute to estimating the positions of the digits relative to one another and the forces generated during object lifting, providing information relevant to performance monitoring, correction, and the updating of sensorimotor memories that guide subsequent lifts. Accordingly, anticipatory control and its learning, position-force coordination, and the magnitude and persistence of anterograde interference provide candidate behavioral readouts for future studies using controlled visuo-proprioceptive discrepancies to test how proprioception contributes to object manipulation.

Prior work suggests that motor behavior in virtual environments depends on whether virtual visual feedback is paired with physical interaction with the target object. In haptic-free virtual grasping, studies have reported longer movement times and changes in the magnitude and timing of maximum grip aperture relative to physical grasping (45). By contrast, reach-to-grasp kinematics more closely resemble real-world performance when participants interact with a physical object while viewing virtual representations of both the hand and object, highlighting the importance of visual context and physical interaction (37). However, this work focused on reach-to-grasp kinematics and did not assess anticipatory force scaling, trial-by-trial coordination between digit position and force, or interference from prior object dynamics.

More direct comparisons with the anticipatory control examined here come from studies of virtual object lifting with device-mediated force feedback (38,46). These studies report slower trial-by-trial learning and differences in force scaling relative to prior real-world findings (8). However, several methodological differences, including the absence of direct within-subject real-world comparisons, make it difficult to isolate the effects of the virtual environment itself. In these studies, the object lifting task was rendered in 3D but displayed on a 2D monitor rather than through an immersive headset, which may alter the interpretation of visual feedback during object interaction (38,46). In addition, both studies required lifting objects with smaller torque magnitudes than those in the real-world comparison dataset, likely reducing the magnitude of error signals, and used a wider range of center-of-mass conditions (38,46), which may increase task complexity and anterograde interference (47,48). Together, these factors make it difficult to attribute slower adaptation and differences in force scaling to virtual environments per se rather than to differences in visual presentation and task structure. By contrast, the present hybrid-VR paradigm combined immersive 3D visual feedback with interaction with a physical object and a direct within-subject comparison to real-world performance, providing a more direct test of whether virtual visual presentation altered object manipulation when physical object interaction was preserved.

In conclusion, no significant differences between hybrid-VR and real-world object manipulation were detected in rapid learning of anticipatory force control, coordinated trial-to-trial variability in digit position and force, or susceptibility to repetition-induced anterograde interference. These findings characterize performance during hybrid-VR object manipulation and provide a baseline for future studies introducing controlled visual-proprioceptive discrepancies to directly examine how visual and proprioceptive information contribute to skilled manual actions.

## Supporting information

https://doi.org/10.5281/zenodo.23961758

## 5 GLOSSARY

CoM: Center of Mass
COP_diff_: Center of Pressure Difference
GF_mean_: Grip Force Mean
Hybrid-VR: Hybrid Virtual Reality
LF_diff_: Lift Force Difference
M_Com_: Compensatory torque
N: Newtons
Nmm: Newton millimeters
VR: Virtual Reality

## 6 ACKNOWLEDGEMENTS

We thank our participants for their contribution to science. We thank Kess Folco and Seiji Koiki for their input on the manuscript.

## 9 AUTHOR CONTRIBUTIONS

C.A.S. and M.M. conceived and designed research; C.A.S. performed experiments; J.Z. rendered the VR environment; C.A.S. and M.M. analyzed data; C.A.S. and M.M. interpreted results of experiments; C.A.S. and M.M. prepared figures; C.A.S. and M.M. drafted manuscript; C.A.S. and M.M. revised manuscript; C.A.S., J.Z., and M.M. approved submission.

## 10 SUPPLEMENTAL MATERIALS

Supplemental Tables: S1-S2: https://doi.org/10.5281/zenodo.23986215

Supplemental Figure: S1: https://doi.org/10.5281/zenodo.23961758

## 12 DATA AVAILABILITY

All data will be made available upon reasonable request. Analysis code is publicly available on our *Motor Skill Lab* GitHub repository (https://github.com/motorskill).

