## Supplementary figures and images for "A behavioral comparison of hybrid virtual reality and real-world object lifting"

### https://doi.org/10.5281/zenodo.23961758

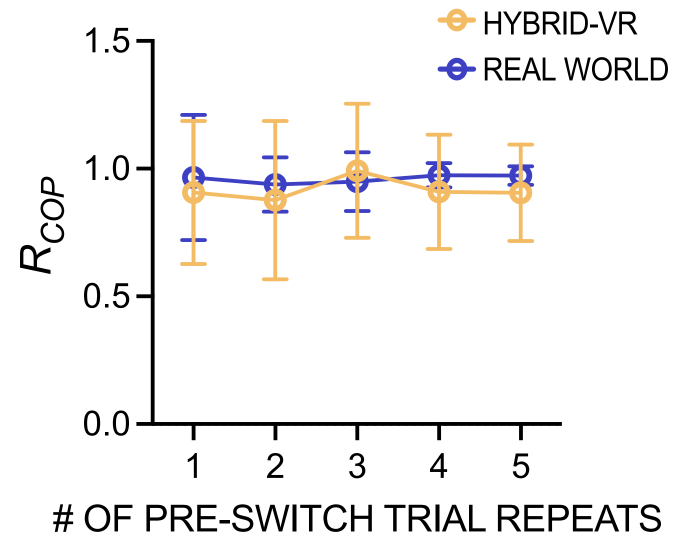
HYBRID-VR
REAL WORLD
1.5
1.0
0.5
0.0
$R_{COP}$
1
2
3
4
5
#### OF PRE-SWITCH TRIAL REPEATS
